# Emotion Modulation of the Body-Selective Areas in the Developing Brain

**DOI:** 10.1101/564633

**Authors:** Paddy Ross, Beatrice de Gelder, Frances Crabbe, Marie-Hélène Grosbras

**Author notes:** Corresponding Author: Paddy Ross Department of Psychology Durham University Science Site South Road Durham DH1 3LE.

## Abstract

Emotions are strongly conveyed by the human body and the ability to recognize emotions from body posture or movement is still developing through childhood and adolescence. To date, there are very few studies exploring how these behavioural observations are paralleled by functional brain development. Furthermore, there are currently no studies exploring the development of emotion modulation in these areas. In the current study, we used functional magnetic resonance imaging (fMRI) to compare the brain activity of 25 children (age 6-11), 18 adolescents (age 12-17) and 26 adults while they passively viewed short videos of angry, happy or neutral body movements. We observed that when viewing bodies generally, adults showed higher activity than children bilaterally in the body-selective areas; namely the extra-striate body area (EBA), fusiform body area (FBA), posterior superior temporal sulcus (pSTS) and amygdala (AMY). Adults also showed higher activity than adolescents, but only in right hemisphere body-selective areas. Crucially, however, we found that there were no age differences in the emotion modulation of activity in these areas. These results indicate, for the first time, that despite activity selective to body perception increasing across childhood and adolescence, emotion modulation of these areas in adult-like from 7 years of age.

**Conflict of Interest:** The author declares that the research was conducted in the absence of any commercial or financial relationships that could be construed as a potential conflict of interest.

## 1 Introduction

A vast literature indicates that the ability to correctly perceive emotions from other people doesn’t reach adult levels until mid-adolescence (Herba, Landau et al. 2006, Chronaki, Hadwin et al. 2015). Moreover, the brain circuits specifically engaged while exposed to a stimulus depicting an emotion also undergo functional and structural changes in the period encompassing late childhood and adolescence (Paus 2005, Blakemore and Choudhury 2006, Lenroot and Giedd 2006, Giedd 2008, Blakemore 2012). For instance, diffusion tensor imaging (DTI) studies have shown that fiber pathways connecting medial and lateral-temporal cortices are reorganized between childhood and adulthood (Loenneker, Klaver et al. 2011). From a functional point of view, event-related potentials (ERPs) studies have shown that the signature of emotion processing when viewing faces doesn’t show adult patterns before the age of 14 (Batty and Taylor 2006). Functional magnetic resonance imaging (fMRI) data in adults also reveals evidence of emotion modulation in the face selective-areas of the visual cortex (fusiform face area, occipital face area, posterior superior temporal sulcus)(Morris, Friston et al. 1998, Rotshtein, Malach et al. 2001, Vuilleumier, Armony et al. 2001, Winston, Vuilleumier et al. 2003, van de Riet, Grezes et al. 2009, Kret, Pichon et al. 2011) but not in children (Thomas, Drevets et al. 2001, Evans, Todd et al. 2010, Todd, Evans et al. 2011), nor in adolescents (Pavuluri, Passarotti et al. 2009, Pfeifer, Masten et al. 2011). Yet, using angry and happy facial expressions, Hoehl, Brauer et al. (2010) found that young children (5-6 years old) displayed heightened amygdala activation in response to emotional faces relative to adults (see also Thomas, Drevets et al. (2001)). Todd, Evans et al. (2011) expanded upon this by demonstrating that children (3.5-8.5 years old), but not adults (18+ years old), showed greater amygdala activation to happy rather than angry faces; although amygdala activation for angry faces increased with age. Monk et al (2003) reported that, when attention is unconstrained adolescents exhibit greater amygdala activity to fearful faces than adults, a finding replicated by Guyer, Monk et al. (2008). For happy and angry faces, developmental differences were observed in the orbitofrontal cortex and anterior cingulate cortex. These data point to a developing emotion recognition system in which age is not simply an additive effect driving increased activation. Rather, the system, including occipitotemporal, limbic and prefrontal regions, shows preferential activation for different emotional expressions that change with age.

### 1.1 Body-Selective Areas

All the aforementioned research has been conducted in relation to the perception of emotion from faces. However, emotion is also strongly conveyed by other means, in particular body posture and movements (Clarke, Bradshaw et al. 2005, de Gelder 2006, de Gelder 2009). Previous work has shown that the capacity to recognize basic emotions from body movements also improves throughout childhood and adolescence (Boone and Cunningham 1998, Lagerlof and Djerf 2009, Ross, Polson et al. 2012). The question of whether related brain processes also change during this time has not yet been addressed.

Several brain areas have been identified as being specialized for the recognition and interpretation of human form and human motion: namely the extra-striate body area (EBA) located bilaterally in the lateral occipitotemporal cortex (LOTC), the fusiform body area (FBA) and areas in the inferior parietal lobe (IPL) and posterior superior temporal sulcus (pSTS) (Downing, Jiang et al. 2001, Peelen and Downing 2005, de Gelder 2006, Weiner and Grill-Spector 2011, Grosbras, Beaton et al. 2012, Basil, Westwater et al. 2016). Research consistently reports modulation by emotion in the fusiform gyrus (de Gelder, Snyder et al. 2004, Grosbras and Paus 2006, de Gelder, Van den Stock et al. 2010, Kret, Pichon et al. 2011) and in the LOTC (Grosbras and Paus 2006, Grèzes, Pichon et al. 2007). Peelen, Atkinson et al. (2007) demonstrated that the strength of emotion modulation in these areas was related, on a voxel by voxel basis, to the degree of body selectivity, but showed no such relationship with the degree of BOLD response when viewing faces. In other words, the emotional signals from the body might ‘modulate’ the complexes of neurons that code for the viewed stimulus (Sugase, Yamane et al. 1999), rather than providing an overall boost in activation for all visual processing in the extra-striate visual cortex. Furthermore, the same authors have shown that the modulation of body-selective areas in adults is also positively correlated with amygdala activation (Peelen, Atkinson et al. 2007). To our knowledge, brain response in children and adolescents when viewing emotional body stimuli has not yet been tested.

### 1.2 Development of Body-Selective Areas

To date, only three fMRI studies have explored the structural and functional development of the body-selective regions in children. While two of them reported that body-selective activity within the EBA and FBA was similar (in terms of location, intensity and extent) in children older than 7 and in adults (Peelen, Glaser et al. 2009, Pelphrey, Lopez et al. 2009), we found that it was not yet adult-like in pre-pubertal children (age 7-11; Ross, de Gelder et al. (2014)). The latter study mirrors findings in the face-selective regions in the ventral stream, which take more than a decade to mature (Scherf, Behrmann et al. 2007, Grill-Spector, Golarai et al. 2008). Of the other two studies, only Peelen, Glaser et al. (2009) included adolescent subjects but did not distinguish them from children in their group analysis, potentially masking any differences between the two age groups. Therefore the question remains open as to how the brain responds to viewing body movements during adolescence. Moreover it is unknown how this brain activity would be modulated by the emotion conveyed by these stimuli. Given that emotional abilities, including emotion perception from social signals, change significantly during adolescence, we might expect different pattern of brain activity in adolescents compared to children and adults.

### 1.3 Current Study

Here, building upon our previous data collected in pre-pubertal participants (Ross, de Gelder et al. 2014), our goal was to investigate, for the first time, the functional development of the EBA, pSTS, FBA and amygdala in relation to the perception of emotional human body movements across childhood and adolescence. Based on the developmental results reported for facial expression perception (Batty and Taylor 2006, Tonks, Williams et al. 2007, Peelen, Glaser et al. 2009, Todd, Evans et al. 2011) we hypothesised that the activity and emotion modulation of the body-selective areas in the visual cortex would develop along a similar protracted trajectory.

## 2. Materials and Methods

### 2.1 Participants

Forty-six primary and secondary school participants were recruited from local schools and after-school clubs in the West End of Glasgow. Three of the younger children were excluded because of excessive head motion in the scanner. Thus data from 25 children (aged 6-11 years: M = 9.55 years, SD 1.46, 13 females) and 18 adolescents (aged 12-17: M = 14.82, SD = 1.88, 9 females) were analysed. Children were all at Tanner stage 1-2, that is, pre-pubertal, as assessed using the Pubertal Developmental Scale (Petersen, Crockett et al. 1988), a sex-specific eight-item self-report measure of physical development (e.g., growth in stature, breast development, pubic hair etc.) filled in by parents. Adolescents were all Tanner stage 3-5; i.e., pubertal or post-pubertal but not yet adults. Binning our subjects in this way avoids arbitrary adolescence age ranges and allows us to have homogenous groups in term of pubertal status. Permission was obtained from managers of after-school clubs and/or head teachers in order to promote the study. Written consent was then obtained from each child’s parent or guardian before testing began (adolescents aged 16 and over were able to provide their own consent). The study was in line with the declaration of Helsinki, was approved by the local ethics board, and all participants understood that participation was voluntary and signed a consent-form. A sample of 26 adult volunteers recruited from the University of Glasgow also took part (aged 18-27 years: M = 21.28 years; SD = 2.11, 15 females). This gave us 69 subjects in total.

### 2.2 Stimuli and Procedure

Forty-five short video-clips were taken from a larger set created and validated by (Kret, Pichon et al. 2011). Each clip depicted one actor, dressed in black against a green background, moving in an angry, happy or neutral manner. Six actors were males and nine females, with each actor recorded three times for each of the three emotions. The videos were recorded using a digital video camera and were edited to two-second long clips (50 frames at 25 frames per second). The faces in the videos were masked with Gaussian filters so that only information from the body was perceived (for full details and validation of stimuli (see Kret et al. (2011), de Gelder and Van den Stock (2011)). In addition, to use as control stimuli, we selected videos depicting non-human moving objects (e.g. windscreen wipers, windmills, metronomes etc.) from the internet. We edited these clips using Adobe Premiere so that they matched the body stimuli in terms of size, resolution, and luminance. A green border matching the colour of the human video background was added. In the fMRI experiment, stimuli were presented in blocks of five clips (10 seconds).

Furthermore, to control for potential low-level parameters effects on fMRI activity, we computed one measure of low-level local visual motion in the clips in order to enter it as a covariate in our fMRI regression analysis. In each clip, we first calculated frame-to-frame change in luminance in the background as a surrogate measure of noise level. Then for each pair of consecutive frames we counted the number of pixels where the change in intensity was higher than noise. We averaged these numbers yielding one value per clip, representing the motion in this clip. Then we computed the cumulative motion for the five clips in each block. Overall the blocks of non-human clips had slightly more motion than the blocks of body movements clips, although this was not statistical significant (*t*(16) = 1.89, *p* = 0.076). In the emotion clips we found a significant difference in movement between the angry and neutral body expressions (*t =* 3.78, *p*<0.005) only. To control for these effects these measures were added a covariate in our fMRI analysis.

#### Data Acquisition

We acquired a series of 246 images of brain activity using a 3T fMRI scanner (Tim Trio, Siemens, Erlangen, Germany) equipped with a 32-channels head coil, using standard EPI sequence for functional scans (TR/TE: 2600ms / 40ms; slice thickness = 3 mm; in plane resolution = 3 x 3 mm). In addition, we performed a high-resolution T1-weighted structural scan (1 mm3 3D MPRAGE sequence) for anatomical localization.

#### Main Experiment

The experiment was programmed with MATLAB using the Psychophysics Toolbox Extensions (Brainard 1997). An experimental run consisted of 48 10-seconds long blocks: eighteen blocks of non-human stimuli (10 seconds; 5 clips), eighteen blocks of human stimuli (three blocks of each emotion) and twelve 10-seconds-long blocks of blank screen as a baseline, in a pseudo-randomized order based on an m-sequence avoiding correlation effects between blocks (Buracas and Boynton 2002). Stimuli were back-projected onto a screen positioned at the back of the scanner bore. Participants were able to view the screen thanks to a mirror attached to the head-coil. They were instructed to maintain their gaze in the centre of the screen.

#### Procedure

Participants were installed comfortably in the scanner. Head motion was restricted by comfortable but tight padding. Parents/guardians were allowed to sit next to their children in the scanning room if they or their child wished (This was the case for three subjects). During the set up and the structural scan they watched a cartoon or movie of their choice. They were first familiarized with the scanner environment and noise with a 3-minutes dummy scan (with the same parameters as the experimental scan), during which they watched a movie. After that we gave them feedback on their head motion and, if head motion was an issue, redid a dummy scan encouraging them to keep still.

Before the main experiment started they were reminded to pay careful attention to the stimuli, to look at the central fixation cross and to keep their head still. Following the scan they were given a short forced-choice emotion recognition task using the same stimuli to gauge their understanding of the emotional content of the stimuli. There was no difference between age groups in emotion recognition accuracy (mean accuracy: 89%, 85% and 89% for children, adolescents and adults respectively; F(2,67)=.787, *p*=.46). Some of the subjects also participated in another independent 8-min functional scan during the same scanning session, before completing the structural scans.

### 2.3 Pre-Processing

We used SPM 8 (Welcome Department of Imaging Neuroscience; www.fil.ion.ucl.ac.uk/spm) to process and analyse the MRI data. The functional data were corrected for motion by using a two-pass procedure. First we estimated the rigid-body transformation necessary to register each image to the first one of the time series and applied this transformation with a 4th Degree B-Spline interpolation. Then we averaged all these transformed images and repeated the procedure to register all images to the mean image. Movement correction was allowed up to 2mm translation or 2 degrees rotation; the three participants who had larger head motion were excluded from the analysis. The realigned functional data were co-registered with the individual 3D T1-weighted scan. First we identified AC-PC landmarks manually and estimated the affine transformation from the mean functional image to the structural image. Then this transformation was applied to the whole realigned time series.

The anatomical scans were then segmented for different tissue types and transformed into MNI-space using non-linear registration. The parameters from this transformation were subsequently applied to the co-registered functional data. Normalising the data from adults, adolescents and children into the same stereotactic template allowed us to directly compare the strength and extent of activation across age groups. Several studies examining the feasibility of this approach have found no significant differences in brain foci locations when the brains of children as young as 6 were transformed to an adult template (Burgund, Kang et al. 2002, Kang, Burgund et al. 2003). These findings, and the careful inspection of the normalized images, gave us confidence that there is no confound of brain size in our results.

Before performing the analyses we smoothed the data using a Gaussian kernel (8mm FWHM). High-pass temporal filtering was applied at a cut off of 128 seconds to remove slow signal drifts.

### 2.4 Whole Brain Analysis

A general linear model was created with one predictor of interest for each of the four conditions (Happy, Angry, Neutral, Non-Body). We added our measure of luminance change (video clip motion) as a covariate, allowing us to control for the increased motion in the anger body movements. The six head-motion parameters were also added as regressors of non-interest. The model was estimated for each participant and we computed the following contrasts of interest between individual parameter estimates: Body>Non-Human; Angry>Neutral; and Happy>Neutral. These contrast images were taken to second-level random effect analysis of variance (ANOVA) to create group-averages separately for children, adolescents and adults and to compare the three groups. For the main group comparison analyses (One-way ANOVA and subsequent post-hoc *t-*tests for each contrast), resulting statistical maps are presented using a threshold of *p*<0.05 after Familywise error (FWE) correction at the voxel level and a cluster extent of a minimum of 10 voxels. Anatomical locations for the peak functional activations were determined with the help of the Harvard-Oxford cortical and sub-cortical structural atlases as implemented in FSLview (Jenkinson, Beckmann et al. 2012). Unthresholded statistical maps are available at http://neurovault.org/collections/4178/.

### 2.5 Region of Interest (ROI) Definition and Analysis

In order to directly address our hypotheses, we focused on six ‘body’ areas identified in Ross, de Gelder et al. (2014) (bilateral EBA, FBA and pSTS) and tested whether their activity was modulated by the emotional content of the body movement across ages, as it is the case in adults (de Gelder, Snyder et al. 2004, Grosbras and Paus 2006, Grèzes, Pichon et al. 2007, Peelen, Atkinson et al. 2007, de Gelder, Van den Stock et al. 2010, Kret, Pichon et al. 2011). ROIs encompassing these areas were created by taking the set of contiguous voxels within a sphere of radius 8mm surrounding the voxel in each anatomical region that showed the highest probability of activation in a meta-analysis of 20 studies examining contrasts between moving body and controls in adults (detailed in Grosbras et al., 2012). We also included ROIs covering the amygdala (AMY) bilaterally in the analysis. These were defined using the WFU PickAtlas software within SPM (Maldjian, Laurienti et al. 2003).

To explore differences in the strength of activity in these ROIs across age groups, we extracted the peak *t*-value from each ROI in each participant for the Body>Non-Human, Angry>Neutral and Happy>Neutral contrasts. We chose the peak *t*-value rather than mean/median ROI response as the peak has been shown to correlate better with evoked scalp electrical potentials than averaged activity (Arthurs and Boniface 2003). Furthermore, the peak is guaranteed to show the best effect of any voxel in the ROI and is unaffected by spatial smoothing and normalization. Lastly, a lowest mean activity in the children and adolescent group will be confounded by the fact that the extent of activation differs across groups (Peelen, Glaser et al. 2009, Ross, de Gelder et al. 2014). Individual peak-t values for each ROI were entered into 3x8 mixed-design ANOVAs, with Age Group and ROI as between and within subject factors respectively. Furthermore, to explore more subtle effects over childhood and adolescence, we also performed further peak *t*-value analyses using age as a continuous variable. This can be found in the Supplementary Material. All statistical analysis was performed using SPSS Version 22.

## 3 Results

### 3.1 Whole brain contrasts

#### 3.1.1 Bodies > Non-Human

##### 3.1.1.1 Within Groups

In adults, viewing dynamic bodies compared to non-human stimuli elicited activation in the right fusiform gyri (including FBA), bilateral pSTS, bilateral occipitotemporal cortices (including EBA), bilateral amygdalae, right inferior frontal gyrus, right precuneus and right precentral gyrus.

Adolescents displayed activation in the bilateral occipitotemporal cortices, bilateral superior temporal sulcus, but at our FWE corrected threshold showed no significant activation in the fusiform gyri or amygdalae.

Children also showed similar activation locations as the adults in the right hemisphere, but showed no significant activation in the left occipitotemporal cortex, left fusiform gyrus, left posterior superior temporal sulcus or left amygdala (Figure 1 and Table 1)

**Table 1.** Regions activated in a whole-brain group-average random-effects analysis contrasting Bodies>Non-Bodies. (*p*<0.05 FWE corrected, cluster extent threshold of 10 voxels, maximum cluster sphere 20mm radius). Coordinates are in MNI space. (OTC=Occipitotemporal Cortex; FG=Fusiform Gyrus; STS= Superior Temporal Sulcus; IFG=Inferior Frontal Gyrus; AMY=Amygdala; Pre=Precuneus; PCG=Precentral Gyrus).

| Region | Adults |  |  |  |  | Adolescents |  |  |  |  | Children |  |  |  |  |
| --- | --- | --- | --- | --- | --- | --- | --- | --- | --- | --- | --- | --- | --- | --- | --- |
|  | x | y | z | t | cm3 | x | y | z | t | cm3 | x | y | z | t | cm3 |
| rOTC | 51 | -73 | 4 | 12 | 7.9 | 51 | -70 | 4 | 7.6 | 3.0 | 48 | -76 | 1 | 8.8 | 4.0 |
| lOTC | -48 | -76 | 7 | 7.7 | 2.7 | -48 | -79 | 4 | 6.2 | 0.8 |  |  |  |  |  |
| rFG | 42 | -46 | -17 | 10 | 2.3 |  |  |  |  |  | 42 | -46 | -17 | 6.0 | 0.4 |
| rpSTS | 51 | -40 | 7 | 9.5 | 15 | 60 | -43 | 13 | 7.5 | 3.3 | 60 | -40 | 13 | 7.3 | 6.1 |
| lpSTS | -63 | -49 | 19 | 7.6 | 3.8 | -54 | -52 | 10 | 6.4 | 2.0 |  |  |  |  |  |
| rIFG | 54 | 20 | 22 | 7.1 | 3.6 |  |  |  |  |  |  |  |  |  |  |
| rAMY | 18 | -7 | -14 | 8.6 | 2.3 |  |  |  |  |  | 18 | -7 | -11 | 6.3 | 0.7 |
| lAMY | -18 | -10 | -14 | 6.3 | 0.4 |  |  |  |  |  |  |  |  |  |  |
| rPre | 3 | -55 | 34 | 5.4 | 0.4 |  |  |  |  |  |  |  |  |  |  |
| rPCG | 48 | 5 | 46 | 7.8 | 3.2 |  |  |  |  |  | 45 | 2 | 46 | 6.4 | 1.2 |

**Figure 1.**
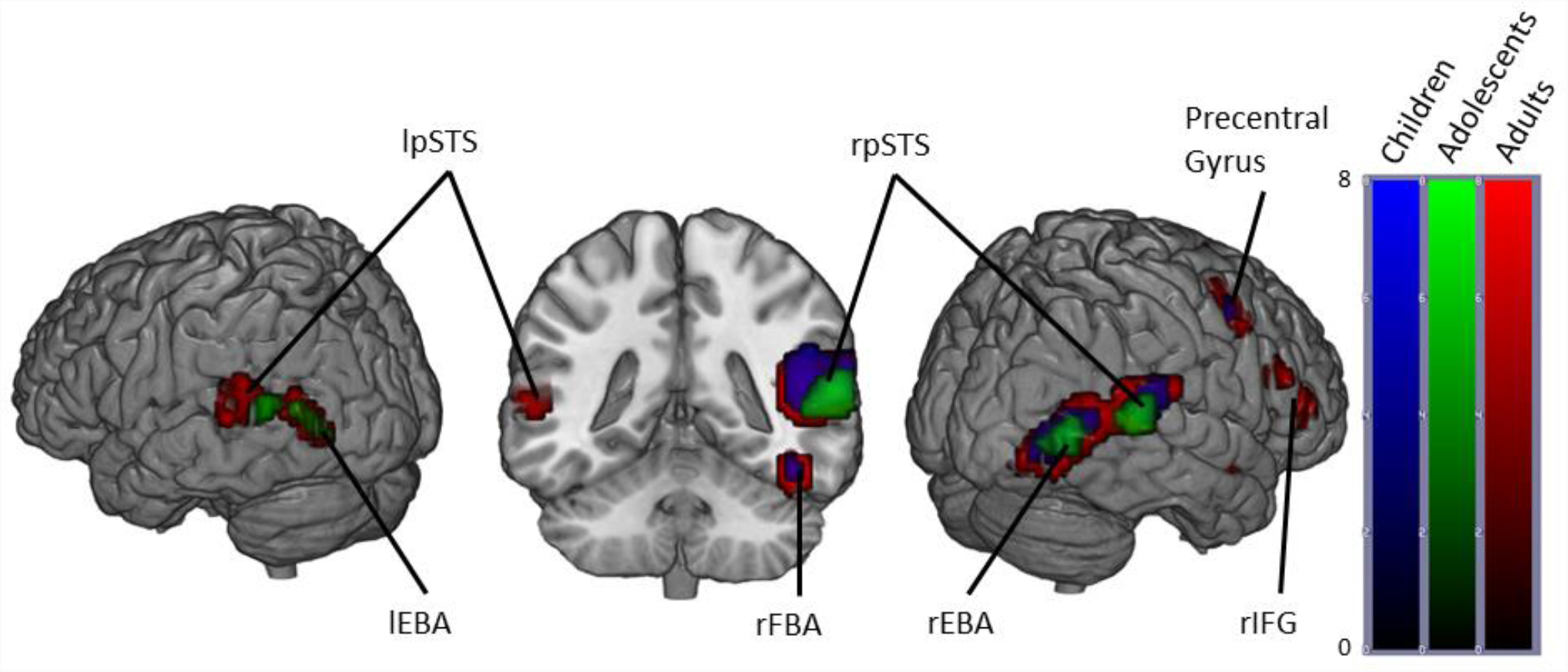
Brain activity when contrasting Bodies > Non-Human stimuli in Children, Adolescents and Adults. (*p*<0.05 FWE corrected, cluster extent threshold of 10 voxels. Colour-bar indicates the threshold of the t-value. Unthresholded statistical maps were uploaded to NeuroVault.org database and are available at http://neurovault.org/collections/4178/.)

##### 3.1.1.2 Between groups

One-way ANOVA of the brain maps of parameters estimates contrasts with Age-group as the between groups factor revealed a main effect of age in the bilateral lingual gyrus (Table 2). We then performed 3 planned comparisons comparing each group individually with each other (Table 3).

**Table 2.** Regions showing a main effect of age in the Bodies>Non-Human contrast. (*p*<0.05 FWE corrected, cluster extent threshold of 10 voxels, maximum cluster sphere 20mm radius). Coordinates are in MNI space.

| Region | Main Effect of Age |  |  |  |  |
| --- | --- | --- | --- | --- | --- |
|  | x | y | z | t | cm3 |
| Left Lingual Gyrus | -9 | -73 | -2 | 19.2 | .57 |
| Right Lingual Gyrus | 9 | -73 | -2 | 17.70 | .27 |

**Table 3.**
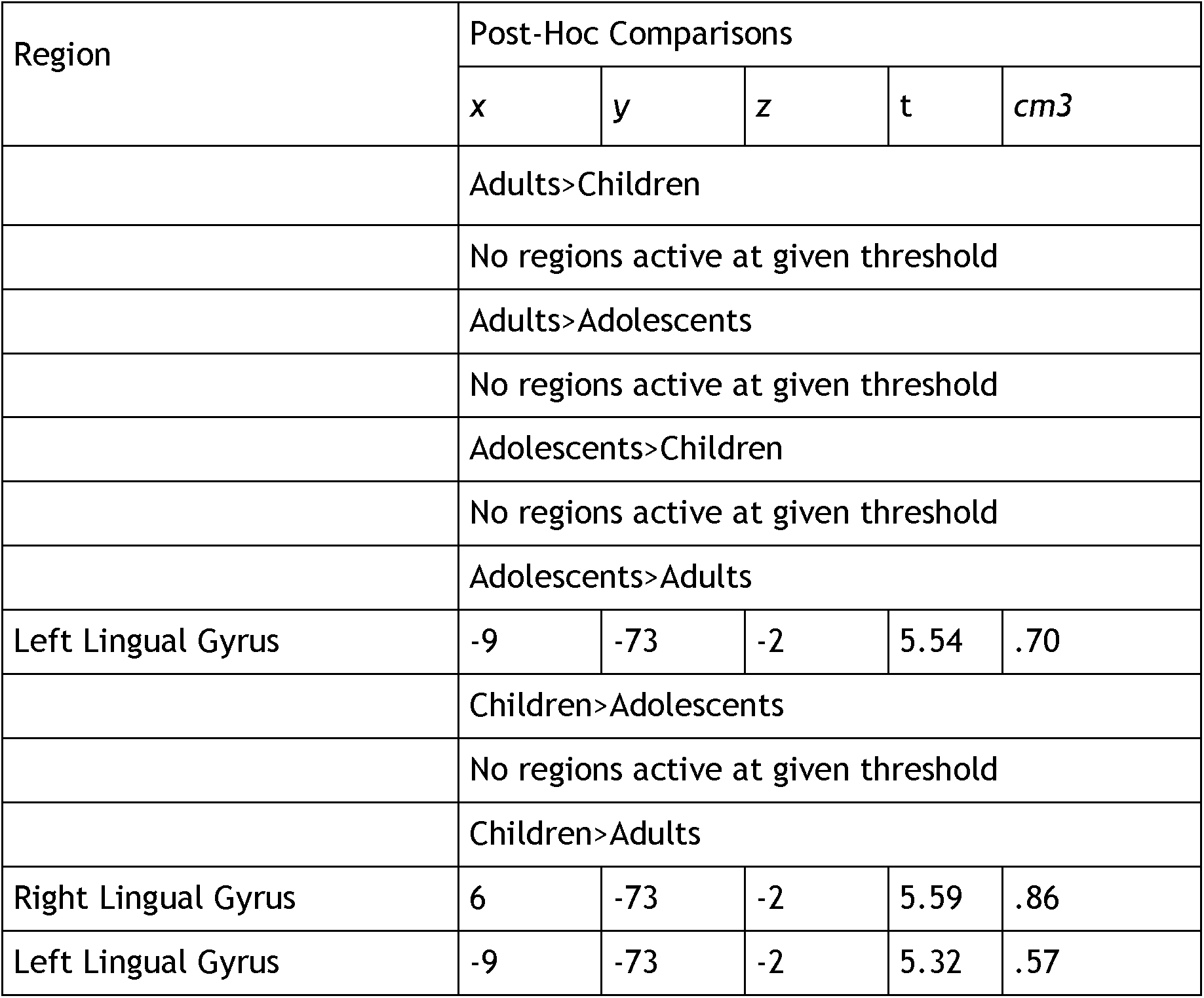
*t*-tests contrasting Bodies>Non-Human across Age contrasts. (*p*<0.05 FWE corrected, cluster extent threshold of 10 voxels, maximum cluster sphere 20mm radius). Coordinates are in MNI space.

| Region | Post-Hoc Comparisons |  |  |  |  |
| --- | --- | --- | --- | --- | --- |
|  | x | y | z | t | cm3 |
|  | Adults>Children |  |  |  |  |
|  | No regions active at given threshold |  |  |  |  |
|  | Adults>Adolescents |  |  |  |  |
|  | No regions active at given threshold |  |  |  |  |
|  | Adolescents>Children |  |  |  |  |
|  | No regions active at given threshold |  |  |  |  |
|  | Adolescents>Adults |  |  |  |  |
| Left Lingual Gyrus | -9 | -73 | -2 | 5.54 | .70 |
|  | Children>Adolescents |  |  |  |  |
|  | No regions active at given threshold |  |  |  |  |
|  | Children>Adults |  |  |  |  |
| Right Lingual Gyrus | 6 | -73 | -2 | 5.59 | .86 |
| Left Lingual Gyrus | -9 | -73 | -2 | 5.32 | .57 |

We found that this main effect was driven by the children and adolescents showing higher bilateral and left lingual gyrus activation than adults respectively.

#### 3.1.2 Angry Bodies > Neutral Bodies

##### 3.1.2.1 Within Groups

When contrasting the conditions Angry bodies and Neutral Bodies, adults showed activation in the bilateral occipital temporal cortices, bilateral occipital fusiform gyri, right middle STS, left posterior STS, right thalamus and right fusiform cortex.

Adolescents displayed activation in the same areas with the exception of the right thalamus.

Children showed activation in the same regions as adults except for the bilateral occipital fusiform gyri. In addition, they showed activation in the left occipital pole, the left superior fusiform gyrus, right hippocampus, left temporal pole, left amygdala and left thalamus (Figure 2 and Table 4).

**Table 4.** Regions activated in a whole-brain group-average random-effects analysis contrasting Angry>Neutral. (*p*<0.05 FWE corrected, cluster extent threshold of 10 voxels, maximum cluster sphere 20mm radius). Coordinates are in MNI space. (OTC=Occipitotemporal Cortex; FG=Fusiform Gyrus; STS= Superior Temporal Sulcus; OFG=Occipital Frontal Gyrus; OP=Occipital Pole; THA=Thalamus).

| Region | Adults |  |  |  |  | Adolescents |  |  |  |  | Children |  |  |  |  |
| --- | --- | --- | --- | --- | --- | --- | --- | --- | --- | --- | --- | --- | --- | --- | --- |
|  | x | y | z | t | cm3 | x | y | z | t | cm3 | x | y | z | t | cm3 |
| lOTC | -48 | -70 | 10 | 7.0 | 2.2 | -51 | -73 | 10 | 6.3 | 0.7 |  |  |  |  |  |
| lpSTS | -51 | -52 | 10 | 4.8 | Sub-Peak |  |  |  |  |  | -63 | -52 | 13 | 5.4 | 0.4 |
| rmSTS | 48 | -43 | 10 | 6.6 | 1.6 |  |  |  |  |  | 63 | -37 | 10 | 6.0 | 2.2 |
| rOTC | 48 | -67 | 7 | 6.2 | 0.9 | 48 | -64 | 7 | 5.5 | 0.68 | 45 | -67 | 7 | 6.2 | 2.7 |
| rTHA |  |  |  |  |  |  |  |  |  |  | 18 | -31 | 1 | 6.2 | 0.7 |
| rFG | 39 | -49 | -17 | 4.5 | 2.19 |  |  |  |  |  | 39 | -49 | -17 | 6.1 | 0.76 |
| rOFG | 27 | -76 | -8 | 5.6 | 0.4 | 18 | -85 | -8 | 6.6 | 1.4 |  |  |  |  |  |
| lOFG | -30 | -76 | -8 | 5.9 | 0.5 |  |  |  |  |  |  |  |  |  |  |
| lOP |  |  |  |  |  |  |  |  |  |  | -9 | -100 | 7 | 5.7 | 1.0 |

**Figure 2.**
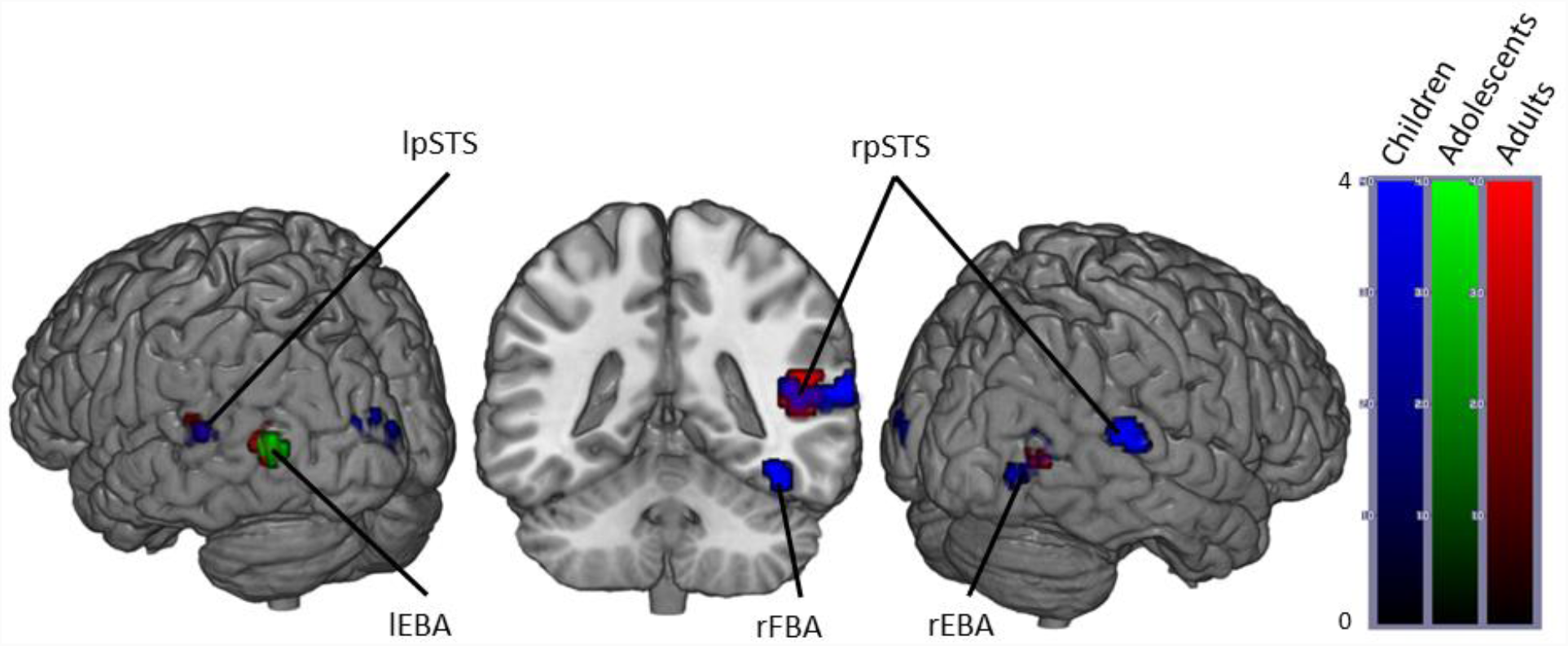
Brain activity when contrasting Angry > Neutral Bodies in Children, Adolescents and Adults.

(*p*<0.05 FWE corrected, cluster extent threshold of 10 voxels. Colour-bar indicates the threshold of the t-value. Unthresholded statistical maps were uploaded to NeuroVault.org database and are available at http://neurovault.org/collections/4178/.)

##### 3.1.2.2 Between Groups

We observed no main effect of Age group when performing a One-Way ANOVA of Angry>Neutral brain maps with Age group as the between subjects factor.

#### 3.1.3 Happy > Neutral

##### 3.1.3.1 Within Groups

The Happy>Neutral contrast in adults revealed activation in the bilateral occipital temporal cortices, right middle and posterior STS and bilateral occipital pole.

Adolescents showed the same pattern of activation except for the right posterior STS. In addition, they also showed activation in the bilateral lingual gyrus and the left posterior STS.

Children only showed significant activation in the right occipitotemporal cortex (Figure 3 and Table 5).

**Table 5.** Regions activated in a whole-brain group-average random-effects analysis contrasting Happy>Neutral. (*p*<0.05 FWE corrected, cluster extent threshold of 10 voxels, maximum cluster sphere 20mm radius). Coordinates are in MNI space. (OTC=Occipitotemporal Cortex; STS= Superior Temporal Sulcus; OP=Occipital Pole; LG=Lingual Gyrus).

| Region | Adults |  |  |  |  | Adolescents |  |  |  |  | Children |  |  |  |  |
| --- | --- | --- | --- | --- | --- | --- | --- | --- | --- | --- | --- | --- | --- | --- | --- |
|  | x | y | z | t | cm3 | x | y | z | t | cm3 | x | y | z | t | cm3 |
| rOTC | 48 | -70 | 4 | 6.8 | 3.7 | 45 | -64 | 7 | 6.4 | 1.2 | 45 | -64 | 7 | 5.1 | 0.5 |
| lOTC | -48 | -73 | 7 | 7.9 | 3.0 | -51 | -73 | 10 | 5.9 | 0.5 |  |  |  |  |  |
| rmSTS | 45 | -40 | 7 | 7.3 | 1.5 | 48 | -37 | 7 | 5.8 | 0.8 |  |  |  |  |  |
| rpSTS | 42 | -58 | 7 | 5.7 | Sub-Peak |  |  |  |  |  |  |  |  |  |  |
| lpSTS |  |  |  |  |  | -54 | -49 | 13 | 5.8 | 0.3 |  |  |  |  |  |
| rLG |  |  |  |  |  | 18 | -85 | -8 | 7.2 | 1.1 |  |  |  |  |  |
| lLG |  |  |  |  |  | -12 | -85 | -11 | 6.0 | 0.8 |  |  |  |  |  |
| rOP | 15 | -85 | -8 | 6.9 | 3.2 | 24 | -91 | 19 | 5.8 | 0.6 |  |  |  |  |  |
| lOP | -12 | -85 | -8 | 8.7 | 3.6 | -18 | -91 | 16 | 5.4 | 0.5 |  |  |  |  |  |

**Figure 3.**
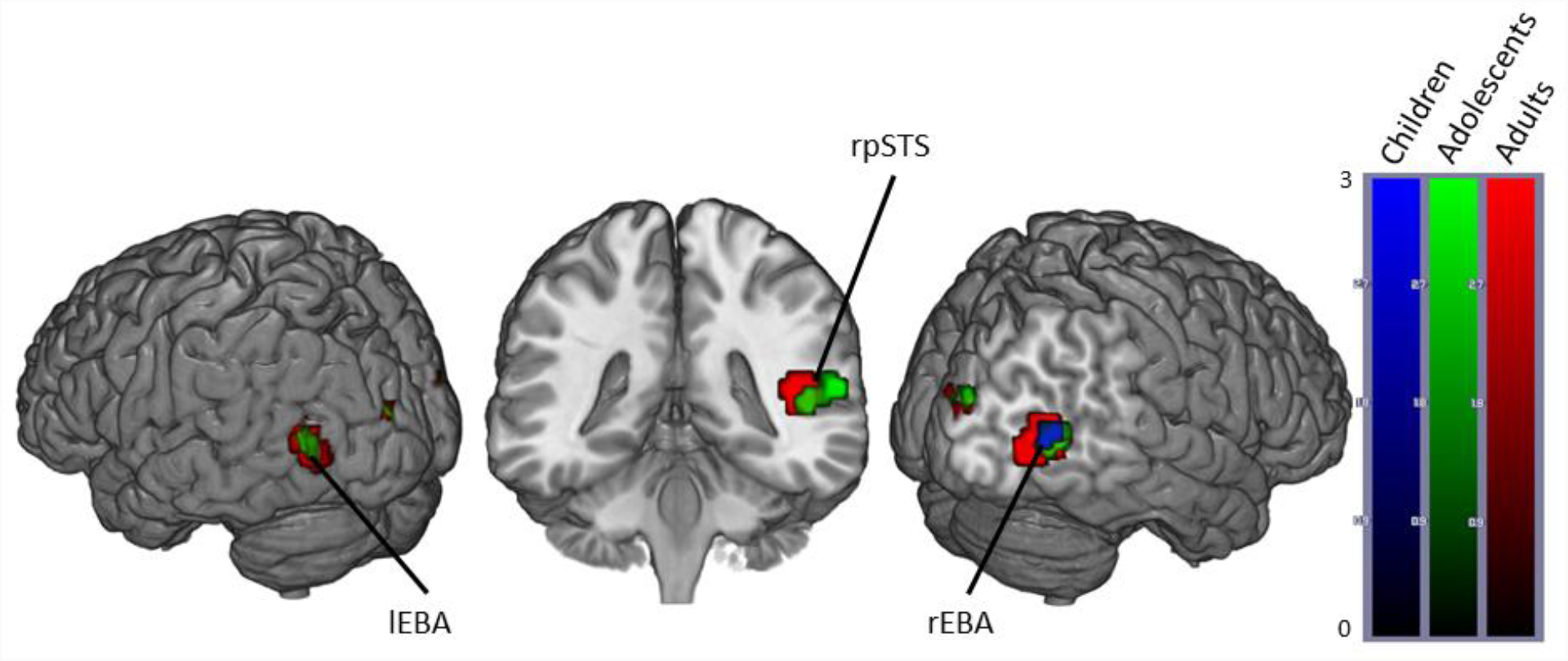
Brain activity when contrasting Happy > Neutral Bodies in Children, Adolescents and Adults. (*p*<0.05 FWE corrected, cluster extent threshold of 10 voxels. Colour-bar indicates the threshold of the t-value. Unthresholded statistical maps were uploaded to NeuroVault.org database and are available at http://neurovault.org/collections/4178/.)

##### 3.1.3.2 Between Groups

We observed no main effect of age when performing a One-Way ANOVA of Happy>Neutral brain maps with Age Group as the between subjects factor.

### 3.2 Region of interest analysis

#### 3.2.1 Bodies > Non-Human

The peak *t*-values for children, adolescents and adults in all eight ROIs under the Bodies > Non-Human contrast are presented in Figure 4. A 3x2x4 Age Group x Hemisphere x ROI ANOVA revealed a main effect of Age Group (F(2,66)=18.72,*p*<.001, η^2^_p_ = .36), driven by adults showing significantly higher peak *t-*values than both children (*p<.*001) and adolescents (*p=*.002). We also found a main effect of ROI (F(3,198)=124.41, *p*<.001, η^2^_p_ = .65) and a main effect of hemisphere (F(1,66)=105.82, *p*<.001, η^2^_p_ = .62) driven by higher peak *t-*values in the right hemisphere (*p<.*001). Further, we observed an interaction between ROI and age group (F(6,198)=4.24, *p*<.001, η^2^_p_ = .11), interaction between hemisphere and age group (F(2,66)=6.48, *p=*.003, η^2^_p_ =.16) but no three-way interaction between hemisphere, ROI and age group (F(6,198)=1.85, *p=*.09, η^2^_p_ = .05).

**Figure 4.**
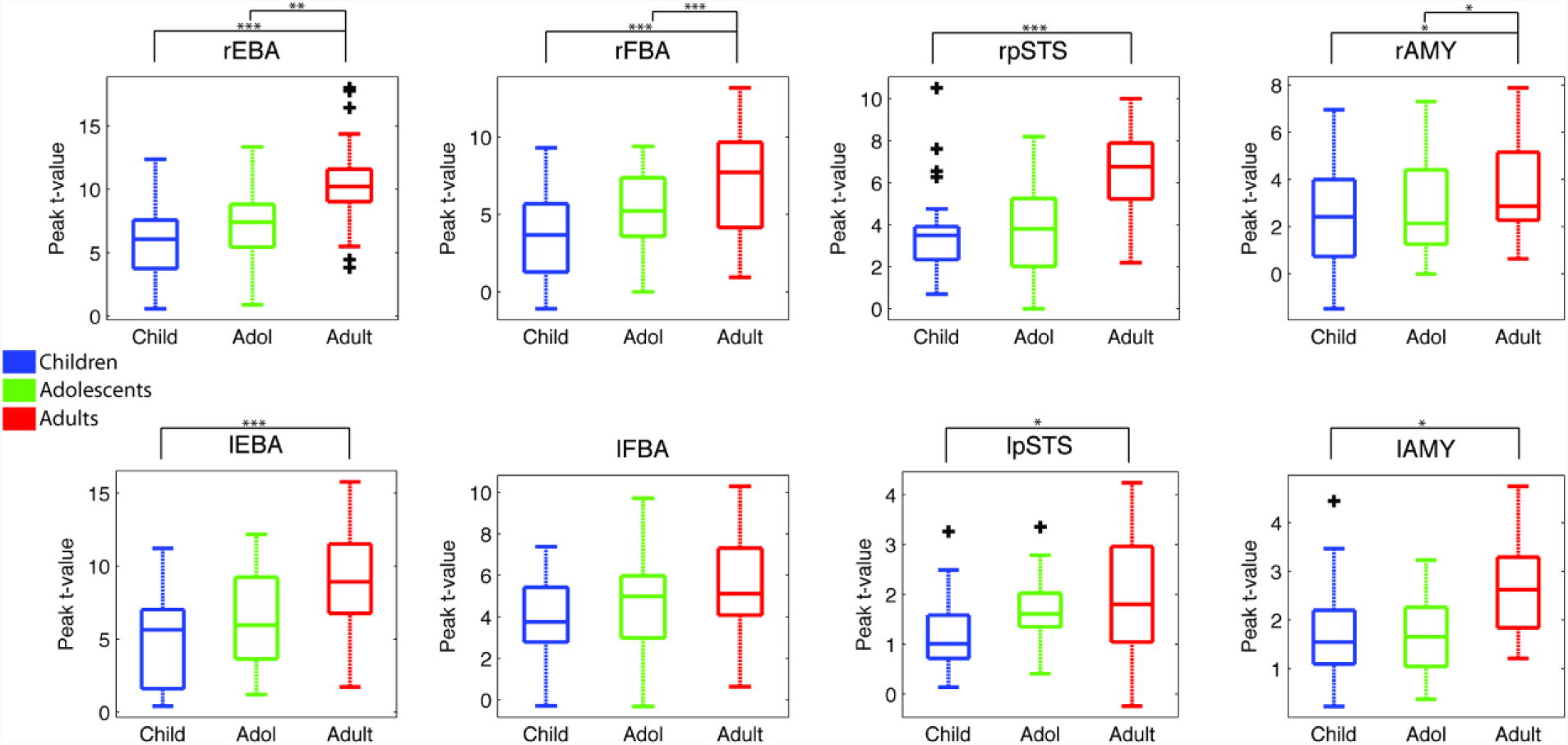
Peak *t*-values in each ROI for each Age Group for the Bodies > Non-Bodies contrast. (It should be noted that the *y-*axis scales are not homogeneous across ROIs). *=*p*<.05; **=*p*<.01; ***=*p*<.001.

Bonferroni corrected follow-up (post-hoc) analysis on the key ROI x age group interaction revealed a main effect of age in all ROIs with the exception of the lFBA (rEBA: F(2,66)=13.83, *p*<.001; lEBA: F(2,66)=10.42, *p*<.001; rFBA: F(2,66)=12.14, *p*<.001; lFBA: F(2,66)=2.06, *p=.*135; rpSTS: F(2,66)=8.55, *p*<.001; lpSTS: F(2,66)=4.00, *p*<.05; rAMY: F(2,66)=5.69, *p*<.005; lAMY: F(2,66)=5.92, *p*<.005).

These effects were driven by adults having increased activity when compared to children in all ROIs except the lFBA, which showed no main effect (rEBA: *p<.*001; lEBA: *p<.*001; rFBA: *p<.*001; lFBA: *p=.*145; rpSTS: *p<.*001; lpSTS: *p<.*05; rAMY: *p<.*05; lAMY: *p<.*05). Adults also showed increased activity over adolescents in the rEBA, rFBA and rAMY (rEBA: *p<.*005; rFBA: *p<.*001; rAMY: *p<.*05). No difference between adolescents and children reached significance.

Exploring the significant hemisphere and age group interaction, post-hoc analysis of hemisphere differences at each age group revealed significantly more right hemisphere activity in all three age groups (Children: *t*(99)=6.7, *p*<.001; adolescents: *t*(71)=5.6, *p*<.001; Adults: *t*(103)=8.4, *p*<.001). We found that there was no significant difference in the right hemisphere between children and adolescents (*p*=.516), but more activity in adults than both children (*p*<.001) and adolescents (*p*<.001). In the left hemisphere there was no significant activity difference between children and adolescents (*p*=.063) or adults and adolescents (*p*>.99) but adults showed significantly more activation than children (*p*<.001).

#### 3.2.2 Emotional Bodies > Neutral Bodies

To explore the developmental trajectories of emotion modulation we broke down the ‘Bodies’ condition into the 3 emotions (Angry, Happy and Neutral). A 3x2x3x4 Age Group x Hemisphere x Emotion x ROI ANOVA of the peak *t-*values revealed the following:

We found a main effect of emotion (F(2,132)=71.03, *p*<.001, η^2^_p_ = .52) which is driven by Angry and Happy giving significantly higher peak t values compared to Neutral (*P*<.001). We found no significant difference between Angry and Happy.

We found a main effect of hemisphere (F(1,66)=79.5, *p*<.001, η^2^_p_ = .55) driven by significantly higher peak t values in the right hemisphere overall.

We found a significant interaction between ROI and emotion (F(6,396)=29.6, *p*<.001, η^2^_p_ = .31) and a significant interaction between ROI and hemisphere (F(3,198)=10.8, *p*<.001, η^2^_p_ = .14).

Crucially, however, none of these effects showed any interaction with age (Emotion x Age F(4,132)=.37, *p*=.83, η^2^_p_ = .01; Hemisphere x Age F(2,66)=2.2, *p*=.123, η^2^_p_ = .06; ROI x Emotion x Age F(12,396)=1.06, *p*=.39, η^2^_p_ = .03; ROI x Hemisphere x Age F(6,198)=1.6, *p*=.136, η^2^_p_ = .05).

Considering Age as a continuous variable, rather than a grouping factor, yielded the same results (presented as Supplementary Material).

## 4 Discussion

We investigated the development of the body-selective areas in the visual cortex and their modulation by emotion by comparing brain activity in adults, adolescents and children passively viewing angry, happy and neutral body movements, as well as objects’ movements. The present results show that adults display bilateral activity in the main body-selective areas of the visual cortex (EBA, FBA and pSTS) when viewing Bodies compared to Non-Human stimuli. For the same contrast, adolescents showed activity in the bilateral EBA and STS, and children in all three areas, but restricted to the right hemisphere. Whole-brain analyses revealed an additional main effect of age-group in the bilateral lingual gyrus driven by higher activation in the children and adolescents compared to adults. Adults showed higher peak *t*-values than children in all the body-selective ROI except the lFBA, and higher peak *t*-values than adolescents in the right regions of interest (rEBA, rFBA and rAMY) except the right pSTS. This result complements our previous work (Ross et al., 2014) while showing that in these regions the magnitude of activity is still lower than adults in adolescents aged 12-17.

In terms of emotion modulation, and contrary to our hypothesis, we did not find any difference between age groups when contrasting Angry or Happy body movements to Neutral body movements: all three age-groups showed similar difference in activation in the body-selective areas and amygdala when viewing emotional compared to non-emotional stimuli.

### 4.1 Body-Selective Areas in Adolescents and Children

This is the first study to compare body-selective areas in children, adolescents and adults. When contrasting viewing Bodies to viewing Non-Human movements, we found similar body-selective areas in children, adolescents, and adults. Interestingly, both children and adolescents showed higher activation in the bilateral lingual gyrus compared to adults. The lingual gyrus has been identified as having a role in various higher-level visual functions such as word processing (Mechelli, Humphreys et al. 2000), complex visual processing (Klaver, Lichtensteiger et al. 2008), and most relevant to this study, the perception of human biological motion (Vaina, Solomon et al. 2001, Servos, Osu et al. 2002, Ptito, Faubert et al. 2003, Santi, Servos et al. 2003). Using diffusion tensor imaging (DTI), (Loenneker, Klaver et al. 2011) found that fiber tracts in the ventral stream differed between adults (aged 20-30) and children (aged 5-7). They found no difference in the fiber tract volume, but instead showed that adults had additional connections to posterior lateral areas (OTC), whereas children showed additional connections to posterior medial areas (LG). In other words, the ventral visual system was ‘adult-like’ in terms of fiber tract volume, but these differences in connection trajectories between children and adults suggests a reorganisation of fiber pathways from medial to lateral-temporal cortex. Thus our data may illustrate developmental differences in processing pathways that could be linked to structural immaturities. Presumably this reorganisation is also not complete by adolescence, as our adolescent group showed the same effect. Loenneker, Klaver et al. (2011) also showed that ventral stream connections to the right fusiform gyrus (containing the FBA) seem to not be completely established in their sample of 5-7 year old children. They suggest that the fiber bundles to the lingual visual areas of the cortex in children may prune until adulthood due to experience-based plasticity. This could explain our finding of increased activity in the lingual gyrus of children and adolescents as compared to adults when viewing body stimuli. Unfortunately their study can shed no light on the functionality of this reorganisation, but it remains plausible then that the fiber tracts in children allow for visual stimuli to reach cortex dedicated to visual memory and language centres. As the child gets older and cerebral architecture is shaped by experience, more fiber bundles develop leading to the OTC, an area specialised in processing visual categories (Grill-Spector 2003).

The ROI analyses revealed higher sensitivity (represented by higher peak *t*-values) in adults than children in all ROIs except the lFBA. The adolescents, however, only showed significantly lower peak *t*-values than the adults in the rEBA, rFBA and rAMY. Given that these three regions also showed a significant difference between adults and children, but no difference between children and adolescents, one could argue for a developmental trajectory that is either so gradual during childhood and adolescence that it is undetectable here, or is simply flat, with maturation occurring at the end of adolescence.

It is notable that these developmental differences occur in the right hemisphere. This may partially support the right lateralisation we previously reported in children viewing body stimuli (Ross et al., 2014), and those observed in children using other modalities such as face stimuli (Golarai, Liberman et al. 2010) or voice processing (Bonte, Frost et al. 2013, Rice, Viscomi et al. 2014). The literature, however, presents a mixed picture with some authors reporting that the cortex becomes less lateralised over age (Golarai, Liberman et al. 2010), increases in lateralisation (Ross et al., 2014), or in the case of Pelphrey, Lopez et al. (2009), a full reversal of lateralisation. Here we find that right lateralisation in our ROIs is present in children and continues through adolescence into adulthood. Altogether, our lateralisation results point towards a stronger activation and later maturation of body-related activity in the right compared to the left hemisphere. This might be related to the change in relative specialization of the different sub-regions of the ventral visual-occipital cortex during literacy maturation and development of the cortex sensitive to words in the left hemisphere (Dehaene, Pegado et al. 2010), although this would need to be tested more specifically.

### 4.2 Emotion Modulation of the Body-Selective Areas

Similar to the Body > Non-Human contrast, the Angry > Neutral and Happy > Neutral contrasts produced activity in the body-selective areas. Interestingly, none of the significant main effects or interactions showed any interaction with age. So, contrary to the increase in activation over age in the Body>Non-Human contrast, the whole-brain Angry>Neutral and Happy>Neutral contrasts and ROI analyses revealed no age differences in emotion modulation of the body selective areas, and no age differences in amygdala response. This rules out an attentional explanation for our Body>Non-Human age differences. If the children were showing significantly lower peak *t*-values due to paying less attention to the stimuli than the adults or adolescents, one would expect that effect to be present in the emotion modulation analysis as well. Furthermore, (Sinke, Sorger et al. 2010) demonstrated that, when presented with socially meaningful stimuli, the body-selective areas were the most active when subjects were not attending the stimulus. Thus, a lack of attention to the stimuli in this study would manifest itself in the data as greater activation (in the context of socially meaningful stimuli).

These findings suggest that even though the body-selective areas are increasing in their levels of recruitment between childhood and adulthood, the emotion modulation of these areas is already adult-like in children. This seems to be in accordance with previous behavioural results into emotion recognition from body movements (Lagerlof and Djerf 2009, Ross, Polson et al. 2012). We previously described a sharp rise in emotion recognition accuracy from full-light human body movements between the ages of 4 and 8.5 years old (Ross, Polson et al. 2012). After 8.5 years we found a much slower rate of improvement in recognition accuracy. In the current study our children subjects range from 6 to 11 years old, so as a group they might be indistinguishable from the adult group in terms of recognition accuracy. In which case finding no age difference in the amygdala response and emotion modulation of the body-selective areas should come as no surprise (indeed, there was no significant age difference in the brief post-scan behavioural emotion recognition task we performed here). It is also possible that the age differences in behavioural performance are linked to the differences in other brain functional circuits (e.g. executive function). It could be the case, as shown using happy and angry faces in Hoehl, Brauer et al. (2010), that 5-6 year old children would show heightened amygdala response to emotional bodies as compared to adults. Indeed, there is evidence that children younger than 6 years of age tend to analyse faces and bodies featurally, whereas older children (like the children in the current study) analyse expressions that include facial and postural cues holistically (Mondloch and Longfield 2010). Exploring these questions in relation to emotional body perception would be a worthwhile extension of the current study; however, brain-imaging data from very young children (4-6 years old) would be needed to explore this possibility.

A question that cannot, by design, be addressed with our data is whether pubertal status could modulate brain response to body stimuli and its modulation by emotion. Indeed, during adolescence chronological age and sexual maturation interact in complex manners on their effect on brain structure and functions (Sisk and Foster 2004, Scherf, Behrmann et al. 2012, Pfeifer and Berkman 2018). In particular, a few studies addressing the development of emotional face processing suggest an effect of puberty even after accounting for age (Forbes, Phillips et al. 2011, Moore, Pfeifer et al. 2012), especially expressed as a reduction in amygdala response to emotion as puberty progresses during mid-adolescence (review in Pfeifer 2018). This effect is very small however, and not observed for all markers of puberty (Goddings, Burnett Heyes et al. 2012). Also, it is not observed in many behavioural studies of basic emotion recognition (Vetter, Leipold et al. 2013, Motta-Mena and Scherf 2017). In light of the current evidence, we do not predict that pubertal status would influence activity related to body perception, and that is why we chose to equate this factor within each age group and focus on age effect. Nevertheless, adequately powered studies for this question could investigate whether puberty and related hormonal changes could have an effect similar to the one suggested for face perception.

Finally, a passive viewing task such as the one employed here cannot address the function of the body-selective system. Using point-light displays, Atkinson, Vuong et al. (2012) showed evidence that emotionally expressive movements do not solely modulate the neuronal populations that code the viewed stimulus category. This implies that a body-selective area that is responding more to emotional compared to neutral stimuli may only be doing so due to top-down influence from some higher cortical area. Pichon, de Gelder et al. (2008) provided evidence, in adults, of amygdala activation as well as activation in the fusiform areas when presented with angry body actions. They attribute this to the brain’s natural response to threat, which has been replicated (van de Riet, Grezes et al. 2009) and mirrored in primate studies (Amaral, Behniea et al. 2003). So, if the stimuli do not modulate the body-selective areas directly, is there any influence from the amygdala? Or, in other words, do the emotional cues contained in the stimuli produce amygdala activity that in turn activates category-specific populations of neurons in the visual cortex? Future work should be designed specifically to test these possibilities.

### 4.3 Conclusions

To summarise, we found evidence that the body-selective areas of the visual cortex are not adult-like bilaterally in children and not adult-like in the right hemisphere in adolescents. Further, we present evidence, for the first time, of emotion modulation in these areas in children and adolescents. We found that emotion modulation of the body-selective areas activity was increased in response to body movements conveying an emotion, positive or negative, compared to neutral movements to the same extent in children and adolescents than in adults. This mirrors various behavioural findings showing that, by the age of 8 years, children recognize emotion from bodily cues with the same proficiency as adults. These data provide new directions for developmental studies focusing on emotion processing from the human body, and could have clinical applications in both typically and atypically developing populations showing deficits in socio-emotional abilities.

## 6 Supplementary Material

### 6.1 Bodies > Non-Human

A 2x4 Hemisphere x ROI ANOVA with Age added as a covariate revealed no main effect of ROI (F(3,201)=1.6, *p*=.2, η^2^_p_ = .02), and no main effect of hemisphere (F(1,67)=0, *p*=.99, η^2^_p_ = .0). Further, we observed no ROI and hemisphere interaction (F(3,201)=1.4, *p=*.24, η^2^_p_ = .02), a significant interaction between ROI and age (F(3,201)=9.5, *p*<.001, η^2^_p_ = .13), interaction between hemisphere and age (F(2,67)=15.75, *p<*.001, η^2^_p_ =.2) and a three-way interaction between hemisphere, ROI and age (F(3,201)=3.9, *p<*.01, η^2^_p_ = .06).

Bonferroni post-hoc analyses revealed that there was a significant increase in activation across age in all ROIs (all *p*<.005) with the exception of the lFBA (*p*=.04; new Bonferroni corrected significance threshold *p*=.00625).

### 6.2 Emotion Bodies > Neutral Bodies

To explore the developmental trajectories of emotion modulation we broke down the ‘Bodies’ condition into the 3 emotions (Angry, Happy and Neutral). A 2x3x4 Hemisphere x Emotion x ROI ANOVA of the peak *t-*values with Age (in months) added as a covariate revealed the following:

We found a main effect of emotion (F(2,134)=8.7, *p*<.001, η^2^_p_ = .11) which was driven by Angry and Happy giving significantly higher peak t values compared to Neutral (*P*<.001). We found no significant difference between Angry and Happy.

We found no main effect of hemisphere (F(1,67)=.56, *p*=.46, η^2^_p_ = .01), nor significant interactions between ROI and emotion (F(6,396)=29.6, *p*<.001, η^2^_p_ = .31) or ROI and hemisphere (F(3,201)=.71, *p*=.55, η^2^_p_ = .01).

Crucially, we also found that none of these effects showed any interaction with age (Emotion x Age F(2,134)=.29, *p*=.75, η^2^_p_ = .00; ROI x Emotion x Age F(6,402)=.98, *p*=.43, η^2^_p_ = .01; ROI x Hemisphere x Age F(2,201)=2.4, *p*=.0.7, η^2^_p_ = .04).

## Acknowledgements

This study was partly funded by the UK ESRC DTC grant ES/J500136/1. MHG was also supported by the AMIDEX foundation (France; grant number A_M-AAC-EM-14-28-140110-16.50).

